# GM-CSF negatively regulates early IL-10 mediated responses

**DOI:** 10.1101/202317

**Authors:** Ruud H. P. Wilbers, Lotte B. Westerhof, Debbie R. van Raaij, Jaap Bakker, Geert Smant, Arjen Schots

**Author notes:** Corresponding author: Ruud H.P. Wilbers, P.O. Box 8123, 6700 ES Wageningen, The Netherlands.

## Abstract

Inflammatory disorders are becoming more prevalent in the Western world. Treatment of these diseases relies on the intervention in inflammatory responses thereby restoring immune homeostasis. One cytokine that has the potential to restore immune homeostasis is the anti-inflammatory cytokine interleukin-10 (IL-10). But until now IL-10 treatment has not been as successful as anticipated. A reason for this may be that IL-10 responsiveness depends on the environment of the inflamed tissue. In this study we describe that granulocyte-macrophage colony-stimulating factor (GM-CSF) is a key cytokine that negatively regulates IL-10-mediated responses. Dendritic cells differentiated from bone marrow with GM-CSF have a reduced ability to respond to IL-10. Dendritic cells are impaired in their up-regulation of IL-10-induced SOCS3 expression and are unable to suppress LPS-induced TNF-α expression at an early time point. Furthermore, GM-CSF treatment partially replicates this phenotype in macrophages. Surprisingly, GM-CSF seems to regulate IL-10 activity in macrophages without affecting STAT3 activation. Still, GM-CSF induces constitutive phosphorylation of glycogen synthase kinase 3β, a signalling component downstream of the PI3K/Akt pathway. Knowledge on the exact mechanism by which GM-CSF negatively regulates IL-10 activity could give novel insights on the integration of signal transduction pathways elicited by different cytokines. Ultimately this knowledge could provide us with new therapeutic strategies to treat inflammatory disorders.

## Introduction

The prevalence of inflammatory disorders, like multiple sclerosis, rheumatoid arthritis, inflammatory bowel disease or allergies, has drastically increased over the last couple of decades. These inflammatory disorders are characterised by uncontrolled immune responses against harmless antigens or commensal bacteria. Treatment of these diseases relies on the intervention in inflammatory responses and thereby restoring immune homeostasis. Many inflammatory disorders are treated with immunosuppressive drugs or monoclonal antibodies that target key pro-inflammatory cytokines, such as tumour necrosis factor alpha (TNF-α). However, immune intervention can also be performed with anti-inflammatory cytokines. One such cytokine capable of restoring homeostasis is interleukin-10 (IL-10). IL-10 is an anti-inflammatory cytokine that suppresses the activity of both antigen presenting cells and lymphocytes and has been considered as a promising therapy for several inflammatory disorders [1]. However, until now IL-10 therapy has not been as successful as previously anticipated.

Systemic administration of recombinant human IL-10 produced in *Escherichia coli* has been studied in phase II clinical trials to treat patients with Crohn′s disease. But, even though IL-10 treatment is well tolerated [2], patients seem to respond differently to IL-10 treatment [3]. Several explanations for why IL-10 treatment has not been a successful therapy for Crohn′s disease have been postulated: (1) systemic administration does not result in an effective dose in the intestine; (2) disease phenotype/severity cause differences in response to IL-10; (3) IL-10 treatment is only successful as preventive therapy; (4) treatment with IL-10 alone is not sufficient; or (5) IL-10′s immunostimulatory effects counterbalance its immunosuppressive effects [4]. To circumvent systemic administration of IL-10, *Lactococcus lactis* was engineered to secrete human IL-10 and used as an oral delivery system [5]. This local delivery system for IL-10 was shown to be successful in two mouse models of intestinal inflammation [5] and was proven to be safe in a small phase I human trial [6]. However, in 2009 a phase II human trial with engineered *L. lactis* in patients with ulcerative colitis was completed, but no significant difference in mucosal healing versus a placebo control was observed (ActoGeniX press release). Future research has to address whether local delivery of IL-10 is indeed a promising therapeutic approach.

Another explanation for IL-10′s lack of efficacy as therapy could arise from differences between patients depending on disease phenotype or severity [4]. For instance, alveolar macrophages have the reduced ability to respond to IL-10 during chronic inflammation [7]. Pre-treatment of these macrophages with TNF-α reduced their ability to respond to IL-10 without affecting IL-10 receptor (IL-10R) expression. Similarly, agonists of Toll-like receptors (TLR) were able to reduce IL-10-mediated phosphorylation and nuclear translocation of the transcription factor STAT3 in macrophages and dendritic cells without lowering surface IL-10R expression [8, 9]. In contrast, IL-10R expression and subsequent IL-10R signalling was reduced in macrophages upon recognition of zymosan [10] or ligation of Fc receptors [11]. Furthermore, cytokines, like IFN-γ, have also been shown to alter IL-10-mediated responses [12]. Altogether, there is quite some evidence that inflammatory mediators can influence IL-10 activity.

Previously we described that bone marrow-derived dendritic cells respond differently to IL-10 when compared to macrophages [13]. Dendritic cells have a reduced capacity to suppress LPS-induced TNF-α, especially when evaluating early IL-10-mediated suppression. The lack of early TNF-α suppression coincided with the impaired ability of dendritic cells to up-regulate SOCS3 expression upon IL-10 treatment. We therefore continued to investigate the underlying mechanism that explains why dendritic cells respond differently to IL-10 than macrophages. In this study we describe that GM-CSF, the cytokine used to differentiate dendritic cells, is a key factor that negatively regulates IL-10 mediated responses. We also show that GM-CSF regulates IL-10 activity without strongly affecting STAT3 activation. In contrast, GM-CSF induces constitutive phosphorylation of GSK-3β, but whether this is the mechanism by which GM-CSF controls IL-10 activity needs further investigation.

## Results

### IL-10 mediated signalling in dendritic cells and macrophages

Previously we observed cell-specific differences in the response to IL-10 in macrophages and dendritic cells [13]. Dendritic cells are strongly impaired in their ability to suppress LPS-induced TNF-α after 2 hours of stimulation, whereas macrophages already inhibit TNF-αexpression by ~75% (Figure 1a). We therefore investigated whether IL-10 signalling differs in these two cell types. First we focused on the activation of the transcription factor STAT3. As shown in figure 1b, IL-10 induces strong tyrosine phosphorylation of STAT3 (Y705) in a dose-dependent manner in both cell types. Upon quantification of relative STAT3 activation, we observed a stronger degree of STAT3^Y705^ phosphorylation in macrophages than in dendritic cells (Figure 1c). However, this was only significantly higher at a dose of 1 ng/ml IL-10. On the other hand, the degree of serine phosphorylation (S727) in macrophages was only significantly higher at a dose of 10 ng/ml (Figure 1d). Yet, when comparing untreated cells with IL-10 treated cells we only observed dose-dependent STAT3^S727^ phosphorylation in macrophages (*P*<0.05, for all concentrations). Untreated dendritic cells already have a higher degree of STAT3^S727^ phosphorylation, which is not further enhanced by IL-10 treatment. Serine phosphorylation of STAT3 could therefore be cell-type specific.

**Figure 1.**
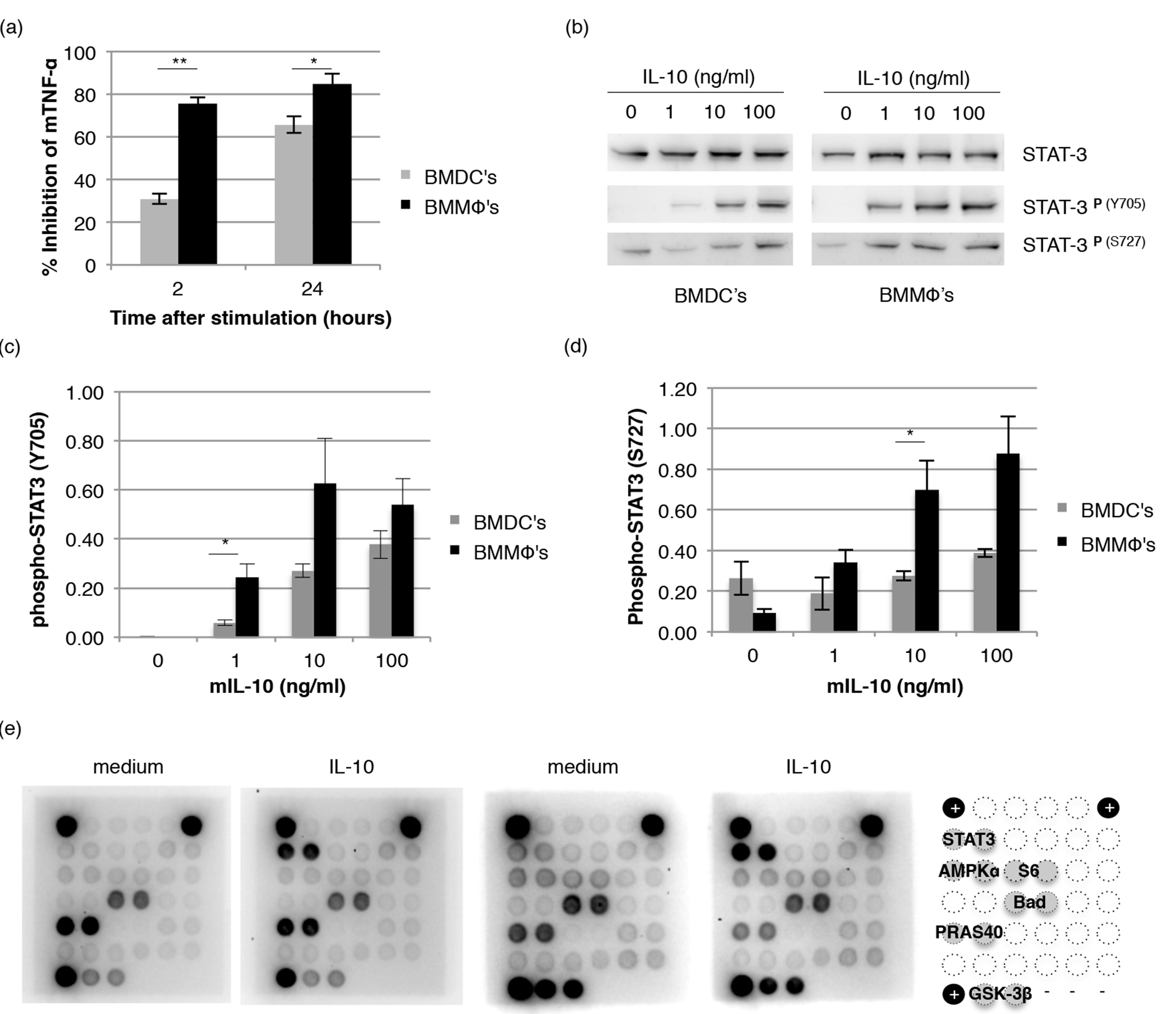
IL-10 mediated signalling in dendritic cells and macrophages. (a) Macrophages and dendritic cells were pre-treated with IL-10 and were stimulated with 100 ng/ml LPS and the inhibition of TNF-α expression was determined at 2 and 24 hours (*n*=3, error bars represent standard error). (b) Phosphorylation of tyrosine 705 (Y705) and serine 727 (S727) of STAT3 by IL-10 (0, 1,10 and 100 ng/ml) was analysed in macrophages and dendritic cells using western blot. (c/d) Relative induction of STAT3 phosphorylation was quantified for Y705 (c) and S727 (d) upon western blot analysis (*n*=3 or 4 for Y705 and S727 respectively, error bars represent standard error). (e) Macrophages and dendritic cells were treated with 100 ng/ml IL-10 and the activation of intracellular signalling pathways was analysed with a PathScan Array. A representative figure is given for 2 independent experiments. Asterisk(s) indicate significant differences as determined by a Welch′s *t*-test (\**P*<0.05; \*\**P*<0.01).

Next, we assessed the activation of 18 well-characterized signalling molecules using the PathScan Intracellular Signalling Array kit. In Figure 1e representative arrays are given for macrophages and dendritic cells treated with 100 ng/ml IL-10 and their respective medium controls. The legend indicates which signalling molecules are differentially affected. Strikingly, we only detected phosphorylation of STAT3^Y705^ upon treatment with IL-10. Interestingly, we did observe differences in the activation of other signalling molecules between untreated macrophages and dendritic cells. Untreated dendritic cells seem to have a higher degree of phosphorylated AMPKα and the ribosomal protein S6, which are indicators for cell cycle progression and cellular growth. Furthermore, several signalling molecules downstream of the PI3K/Akt pathway (PRAS40, Bad and GSK-3β) are activated in untreated macrophages and dendritic cells, but the degree of GSK-3β phosphorylation (Ser9) is much stronger in dendritic cells. Phosphorylation of GSK-3β inhibits its activity and thereby promotes cell survival. Altogether we conclude that STAT3 is the major downstream signalling molecule for IL-10. Furthermore, macrophages and dendritic are differentially affected in signalling molecules that regulate cellular growth and survival.

### GM-CSF negatively regulates IL-10 activity

Dendritic cells are differentiated from bone marrow cells by culturing them in the presence of the cytokine GM-CSF. In order to find out if GM-CSF could be responsible for the altered responses of dendritic cells to IL-10, we investigated whether GM-CSF could replicate this phenotype in macrophages. Macrophages and dendritic cells were differentiated from bone marrow, but now GM-CSF was added to macrophages or depleted from dendritic cells for the last 24 hours of culture. Cells were then pre-treated with IL-10 and subsequently challenged with lipopolysaccharide (LPS). Figures 2a-c reveals that GM-CSF is indeed able to alter the response of macrophages towards IL-10. IL-10 inhibits TNF-α expression in a dose-dependent matter in macrophages independent of GM-CSF treatment (Figure 2a), but GM-CSF treatment lowers the maximum percentage of TNF-α expression at a lower dose of IL-1 0.

**Figure 2.**
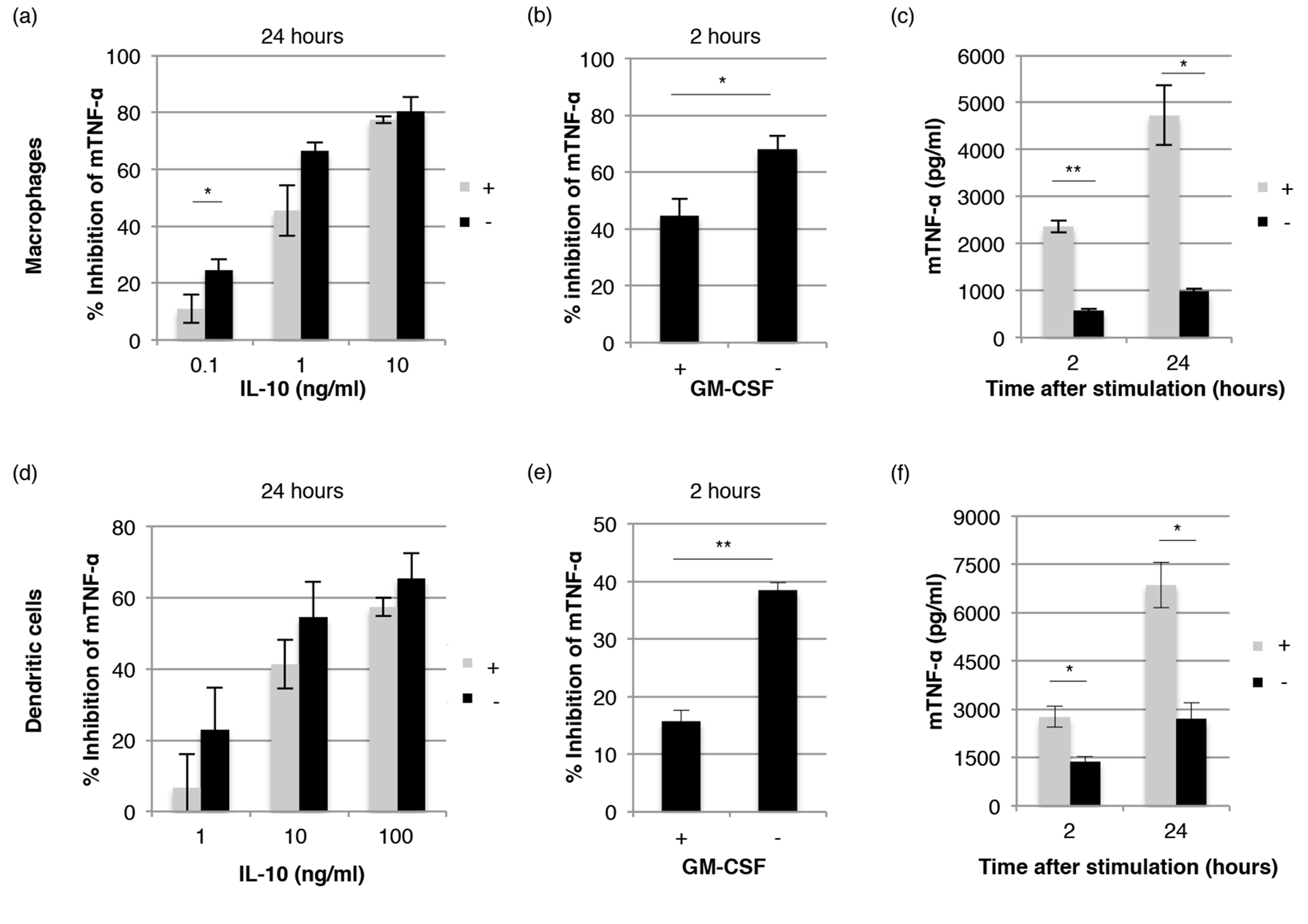
GM-CSF suppresses early IL-10-mediated responses. Macrophages (a-c) and dendritic cells (d-f) were cultured for the last 24 hours in the presence or absence (+/-) of GM-CSF. Cells were pre-treated for 20 min. with IL-10 followed by stimulation with 100 ng/ml LPS. The inhibition of TNF-α expression was determined after 24 hours of stimulation (a/d). Alternatively, cells were pre-treated for 20 min with 10 ng/ml IL-10 and then stimulated with 100 ng/ml LPS. The inhibition of TNF-α expression was determined after 2 hours of stimulation (b/e). Absolute expression levels of TNF-α for the abovementioned experiments are also given (c/f). All bars represent the average of 3-5 biological replicates (*n*=3-5, error bars indicate standard error) and asterisk(s) indicate significant differences as determined by a Welch’s *t*-test (\**P*<0.05; \*\**P*<0.01).

Furthermore, GM-CSF treatment of macrophages significantly reduced early IL-10-mediated suppression of TNF-α by ~25%. We also observed that macrophages treated with GM-CSF produced significantly more TNF-α than untreated macrophages (Figure 2c). These results indicate that GM-CSF is able to alter IL-10-mediated responses in macrophages as was observed for GM-CSF differentiated dendritic cells.

Similarly, depletion of dendritic cells from GM-CSF enhanced their response towards IL-10. No differences were observed in the suppression of TNF-α expression after 24 hour stimulation when dendritic cells were depleted from GM-CSF (Figure 2d). However, early IL-10-mediated suppression of TNF-α expression increased significantly with almost 25% when dendritic cells were depleted from GM-CSF. Also, production of TNF-α was significantly reduced when dendritic cells were cultured in the absence of GM-CSF. We therefore conclude that GM-CSF is a key factor that modulates early IL-10-mediated responses in both macrophages and dendritic cells.

### GM-CSF regulates IL-10 activity without affecting STAT3 activation

Early IL-10 mediated responses seem to be controlled by the major anti-inflammatory pathway Jak1-STAT3-SOCS3. To investigate how GM-CSF influences early IL-10-mediated responses we first focused on the activation of STAT3. As shown in figure 3a, IL-10 induces strong STAT3^Y705^ phosphorylation regardless of GM-CSF treatment. Phosphorylation of STAT3^S727^ seemed also not to be affected by GM-CSF treatment. GM-CSF seems to regulate early IL-10-mediated responses without affecting STAT3 activation. However, early inhibition of TNF-α by IL-10 is completely abrogated in GM-CSF treated macrophages from LysMcre/Stat3flox mice that lack Stat 3 expression in macrophages (Figure S1). This indicates that STAT3 is required for early IL-10-mediated responses, but that GM-CSF negatively regulates this response in a STAT3 independent manner.

As STAT3 phosphorylation was not affected by GM-CSF we continued to investigate the effect of GM-CSF on IL-10-induced SOCS3 expression. Relative SOCS3 transcript levels were determined by quantitative PCR in both macrophages and dendritic cells. As shown in figure 3b, IL-10 induces SOCS3 expression by ~100-fold in macrophages, whereas SOCS3 up-regulation is significantly lower in dendritic cells (12-fold, *P*=0.008). Unexpectedly, no significant differences were found for relative SOCS3 expression levels upon IL-10 treatment of normal and GM-CSF treated macrophages. However, the IL-10 specific induction of SOCS3 was reduced ~10-fold. This is because GM-CSF itself already enhances SOCS3 expression in macrophages prior to IL-10 treatment. Depletion of GM-CSF from dendritic cells only resulted in a 2-fold increase in SOCS3 expression. Therefore GM-CSF seems to regulate early IL-10-mediated responses not only in a STAT3 independent manner, but also without affecting SOCS3 expression levels.

Next, we assessed whether GM-CSF treatment of macrophages alters IL-10-mediated signalling or other signalling pathways by using the PathScan Intracellular Signalling Array kit. As we observed previously, IL-10 only induces STAT3^Y705^ phosphorylation in macrophages and dendritic cells, which is independent of GM-CSF pre-treatment. GM-CSF itself induces an activation state in macrophages that is indistinguishable from dendritic cells. GM-CSF treatment of macrophages enhances the phosphorylation of AMPKα and the ribosomal protein S6, but most strongly induces the phosphorylation of GSK-3β. Phosphorylation of other signalling molecules downstream of PI3K/Akt (PRAS40 and Bad) is also enhanced. On the other hand, depletion of dendritic cells from GM-CSF relieves PRAS40 from phosphorylation, whereas GSK-3β remains strongly phosphorylated. We therefore conclude that GM-CSF alters the activation status of signalling molecules that regulate cellular growth and survival in macrophages and dendritic cells.

**Figure 3.**
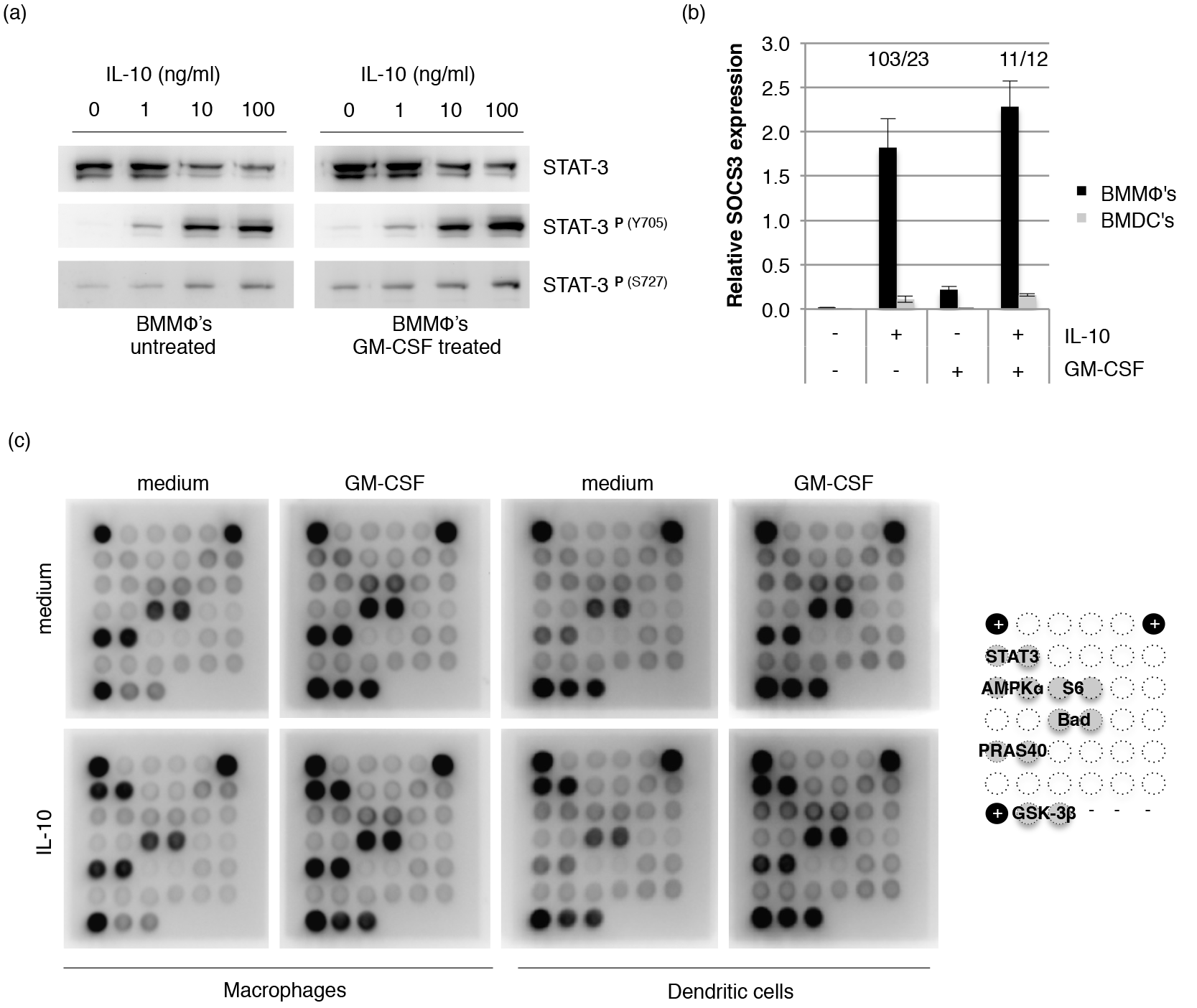
GM-CSF negatively influences IL-10 activity without affecting STAT3 activation. (a) Macrophages were treated for 24 hours with GM-CSF or left untreated. Cells were then activated with IL-10 (0, 1, 10 and 100 ng/ml) and phosphorylation of tyrosine 705 (Y705) and serine 727 (S727) of STAT3 was analysed by western blot. (b) Relative SOCS3 transcript levels were analysed by quantitative PCR in both macrophages and dendritic cells that were treated overnight by GM-CSF or were left untreated (*n*=3, error bars represent standard error). Fold induction of SOCS3 expression upon IL-10 treatment (100 ng/ml) was calculated using the 2^ΔCt^ method using HPRT as a reference gene and is depicted above the bars in the graph (macrophages/dendritic cells). (c) Macrophages and dendritic cells were treated overnight with GM-CSF or left untreated. Cells were then stimulated with 100 ng/ml IL-10 and the activation of intracellular signalling pathways was analysed by PathScan Array. A representative figure is given for 2 independent experiments. Asterisk(s) indicate significant differences as determined by a Welch′s *t*-test (\**P*<0.05; \*\**P*<0.01).

## Discussion

Interleukin-10 (IL-10) is an anti-inflammatory cytokine with promising therapeutic potential, but to date IL-10 therapy has not been as successful in the clinic as previously anticipated. A likely explanation for this phenomenon is that IL-10 activity is influenced by the cytokine milieu present in inflamed tissues [10–12], Previously we have described the differential response of macrophages and dendritic cells towards IL-10 [13]. Dendritic cells are unable to respond rapidly to IL-10 as they are unable to suppress LPS-induced TNF-α at early stages. Furthermore, IL-10-induced SOCS3 expression was strongly reduced compared to macrophages. In this study we further investigated the mechanism that controls this differential response between macrophages and dendritic cells. Our study demonstrates that the cytokine GM-CSF, the cytokine used to differentiate dendritic cells, is a key factor that negatively regulates IL-10 activity. GM-CSF pre-treatment of bone marrow-derived macrophages also reduced their ability to suppress LPS-induced TNF-α at 2 hours after stimulation. Vice versa, depletion of dendritic cells from GM-CSF partially restored their early response to IL-10. Furthermore, both macrophages and dendritic cells cultured in the presence of GM-CSF produced significantly higher levels of TNF-α, which coincided with previous reports from Fleetwood and co-workers [14, 15]. Increased TNF-α expression levels upon LPS stimulation are caused by increased basal TNF-α mRNA transcript levels upon GM-CSF treatment [15]. In our study GM-CSF treated macrophages and dendritic cells do not produce significantly different levels of TNF-α, whereas both cell types respond significantly different towards IL-10. Differences in basal TNF-α expression levels cannot explain the observed differential response towards IL-10. Therefore, we continued to investigate IL-10-mediated signalling in macrophages and dendritic cells.

IL-10 receptor engagement results in the activation of downstream janus kinases Jak1 and Tyk2, which in turn phosphorylate the transcription factors STAT1, STAT3 and in some cell types STAT5 [16–18]. IL-10 is well known for its ability to activate the Jak- STAT pathway [19], but alternative signalling pathways have been described as well. Especially IL-10-mediated activation of the PI3K signalling pathway has been implicated in IL-10′s survival promoting properties [20, 21]. Zhou and co-workers also revealed that IL-10 induced the activation of extracellular signal-related kinase [22]1/2 in a PI3K-dependent manner in promyeloid cells [21]. Furthermore, IL-10 has been shown to activate p38 MAPK signalling to induce the expression of heme oxygenase-1 [23]. In our study we could only detect IL-10-mediated activation of the transcription factor STAT3 in bone marrow-derived macrophages and dendritic cells. This indicates that STAT3 is the major transcription factor required for IL-10-mediated responses in bone marrow-derived cells. However, we did observe subtle differences in the phosphorylation of STAT3^Y705^ between regular macrophages and dendritic cells. More striking was the observation that IL-10 was not able to induce STAT3^S727^ phosphorylation in a dose-dependent manner in dendritic cells. Serine phosphorylation of STAT3 has been reported previously for IL-6, IL-22, IFN-y and EGF [24–27] and enhances transcriptional activity of STAT3. The lack of STAT3^S727^ phosphorylation might explain why IL-10-induced SOCS3 expression and early inhibition of TNF-α expression are impaired in dendritic cells.

Qasimi and co-workers previously described that SOCS3 is required by IL-10 to suppress TNF-α expression at early time points of stimulation [28]. We therefore investigated the role of SOCS3 in more detail. We observed a reduction in the ability of macrophages to respond rapidly to IL-10 upon GM-CSF treatment, but this seemed to be independent from upregulation of SOCS3 expression. Relative induction of SOCS3 expression by IL-10 was hardly affected by overnight treatment of macrophages with GM-CSF or depletion of dendritic cells from GM-CSF. This is in contrast to the difference in IL-10-induced SOCS3 expression between regular macrophages and dendritic cells. Furthermore, we also did not observe a difference in the phosphorylation of STAT3^Y705^ and STAT3^S727^ between regular macrophages or GM-CSF treated macrophages. Therefore, GM-CSF seems to negatively regulate early IL-10-mediated responses in macrophages independent from STAT3-induced SOCS3 expression.

Interestingly, we did find major differences in the activation status of signalling molecules downstream of PI3K/Akt signalling in macrophages and dendritic cells. Furthermore, GM-CSF induced an activation state in macrophages that was indistinguishable from dendritic cells. Particularly the strong serine phosphorylation of GSK-3β^S9^ was a striking observation in GM-CSF cultured dendritic cells or GM-CSF treated macrophages. The degree of GSK-3β^S9^ phosphorylation also seems inversely correlated with early IL-10-mediated responses. A higher degree of GSK-3β^S9^ phosphorylation results in reduced suppression of LPS-induced TNF-α expression by IL-10. As IL-10 is also able to trigger the PI3K signalling pathway it is of great interest to investigate how these signalling pathways of IL-10 and GM-CSF are integrated and result in impaired IL-10-mediated responses.

Activation of STAT3 is regulated by a diverse set of post-translational modifications, including phosphorylation, acetylation and methylation [26, 27, 29–34]. Every modification plays its own role in regulating optimal STAT3 dimerization, DNA binding activity and transcriptional activity [35]. Involvement of the PI3K signalling pathway in the activation of STAT3 has also been reported previously. Spencer and co-workers revealed that the cytomegalovirus homologue of IL-10 was capable of inducing serine phosphorylation of STAT3^S727^ in a PI3K-dependent manner [36]. Furthermore, acetylation of STAT3^L685^ by IL-6 was also shown to depend on PI3K/Akt activation [37]. However, whether IL-10-induced STAT3 transcriptional activation requires acetylation and/or methylation and whether the PI3K pathway is involved, needs further investigation. Interestingly, Waitkus and co-workers recently reported on a novel mechanism of activation of STAT3, which was mediated by GSK-3α/β [35]. GSK-3α/β was shown to directly phosphorylate STAT3 at the residues S727 and T714. The requirement of GSK-3α/β in IL-10-induced STAT3 activation is therefore of great interest as this study shows that GM-CSF strongly inhibits GSK-3β by phosphorylation of serine residue 9. GSK-3α/β might therefore be a key signalling node that is able to control IL-10-mediated responses.

Previously, Hart and co-workers already reported that IL-10 was unable to suppress MHC-II expression in GM-CSF cultured monocytes [38]. Our study now also demonstrates that GM-CSF is able to alter IL-10-mediated suppression of TNF-α expression in both macrophages and dendritic cells. Furthermore, we and others show that GM-CSF cultured cells produce significantly higher levels of the pro-inflammatory cytokine TNF-α [15], but also the secretion of IL-12p70 and IL-23 is significantly enhanced by GM-CSF treated macrophages [15]. IL-23 has recently been identified as an important pro-inflammatory cytokine driving both innate and T cell-induced intestinal inflammation [39]. Therefore, GM-CSF seems to induce a cytokine environment favourable for chronic inflammation. Over the last couple of years GM-CSF has been identified as a key contributor to the development of chronic inflammation in animal models of intestinal inflammation, multiple sclerosis and rheumatoid arthritis [40–42]. Taking together the key role of GM-CSF in the development of chronic inflammation with the results obtained in this study might explain why IL-10 therapy has not been as effective as previously anticipated. Our study demonstrates that the pro-inflammatory cytokine GM-CSF is able to negatively regulate IL-10-mediated responses in macrophages and dendritic cells. However, future research has to elucidate the exact mechanism on how GM-CSF and IL-10 signalling pathways are integrated and result in impaired cellular responses towards IL-10. Ultimately this knowledge could provide us with new therapeutic strategies to treat inflammatory disorders.

## Experimental procedures

### Mice

Wild-type C57BL/6J mice were bred and maintained under specific pathogen-free conditions in the animal facilities at Wageningen University. All experiments were approved by and conducted in accordance with relevant guidelines and regulations of the institutional animal care body at Wageningen University. The experiments of this specific study were approved by the animal experiments committee (DEC) of Wageningen University & Research.

### Bone marrow cultures

Bone marrow was isolated from the femur and tibia of 6-12 week old C57BL/6J mice. Bone marrow derived macrophages (BMMΦ′s) were differentiated at 37°C/5% CO_2_in RPMI-1640 medium containing 4 mM L-glutamine, 25 mM HEPES and supplemented with 10% fetal calf serum, 50 μM β-mercaptoethanol, 50 U/ml penicillin and 50 μg/ml streptomycin and 20% spent medium from L929 cells (ATCC). Bone marrow cells were seeded at 1×10^6^ cells/ml in 6- or 96-well tissue culture plates and cultured for 6 days, while refreshing medium at day 3. After 6 days of culture ≥95% of the cells expressed the macrophage markers F4/80 and CD11b.

Bone marrow derived dendritic cells (BMDC's) were differentiated for 10 days as described [43] using 10% spent medium from murine GM-CSF transfected X63 cells [44]. X63-GM-CSF cells were kindly provided by dr. M. Lutz (University of Erlangen-Nuremberg) with approval of dr. B. Stockinger (MRC National Institute for Medical Research). Briefly, bone marrow cells were plated at 2×10^5^ cells/ml in bacteriological petri dishes and incubated at 37°C/5% CO_2_. At day 3, 6 and 8 medium was refreshed and at day 10 both adherent and non-adherent cells were harvested. At this time typically ~90%of the cells expressed the dendritic cell markers CD11c and MHC class II.

### Flow cytometry

Bone marrow derived cells were stained in FACS buffer (PBS containing 0.1% BSA and 5mM EDTA) using the following monoclonal antibodies for cell surface markers (all obtained from eBioscience): PE-conjugated anti-CD11b, APC-conjugated anti-F4/80, PE-conjugated anti-CD11c, APC-conjugated MHC-II. Cells were first incubated with Fc receptor block (eBioscience) for 10 min to block any non-specific binding and subsequent staining steps were performed for 20 min. at 4°C, followed by washing with FACS buffer. Stained cells were acquired using a Cyan-ADP Analyzer (Beckman Coulter) and analysed with FlowJo software (Tree Star, Inc.).

### LPS stimulation assays

BMMΦ′s were differentiated in 96 well plates and BMDC′s were seeded in 96 well plates at a density of 5x10^4^/well. Cells were pre-treated for 15 min. with IL-10 and subsequently stimulated with 100 ng/ml lipopolysaccharide (LPS). After 2 hours or overnight stimulation, supernatants were analysed for TNF-α using the Ready-Set-Go!^®^ ELISA kit (eBioscience) according to the supplier's protocol.

### IL-10 induced signalling

Bone marrow-derived cells were treated for 20 min. with IL-10 (0, 1, 10 or 100 ng/ml). Cells were lysed using 1x Cell Lysis Buffer (Cell Signaling Technology) and total soluble protein content in the lysates was determined by the BCA method (Pierce). Proteins were separated on 12% Bis-Tris gels followed by transfer to a PVDF membrane by a wet blotting procedure. Thereafter the membrane was blocked in PBS (containing 0.1% v/v Tween-20 and 5% w/v non-fat dry milk powder) for 1 hour at room temperature, followed by overnight incubation at 4°C with monoclonal antibodies specific for STAT3, phospho-STAT3^Y705^ or phospho-STAT3^S727^ in PBS (containing 0.1% v/v Tween-20 and 1% w/v BSA). All STAT3 antibodies were obtained from Cell Signaling Technology. A HRP-conjugated donkey anti-rabbit IgG (Jackson ImmunoResearch) was used as a secondary antibody. Band intensities were analysed with Image J Software.

Cell lysates were also used to investigate the activation of 18 well-characterized signalling molecules using the PathScan Intracellular Signaling Array kit (Cell Signaling Technology) according to the manufacturer′s protocol. Blots and arrays were visualised in the G:BOX Chemi System (Syngene).

### Quantitative PCR

BMMΦ′s were differentiated in 6 well plates and 3×10^6^ BMDC′s were seeded in 6 well plates and were treated for 2 hours with IL-10 (0, 1, 10 or 100 ng/ml). Cells were then washed with PBS and mRNA was isolated with the Maxwell^®^ 16 Tissue LEV Total RNA Purification Kit and the Maxwell^®^ 16 instrument (both from Promega). Then cDNA was synthesized using the GoScript^TM^− Reverse Transcription System (Promega) according to the supplier′s protocol. Samples were analysed in triplicate for SOCS3 and HPRT (reference gene) expression by quantitative PCR using ABsolute SYBR Green Fluorescein mix (Thermo Scientific). Fold induction of SOCS3 expression was determined by the Pfaffl method [45].

### Data analysis

All data shown in the figures indicate the average of at least three biological replicates (*n*) that were determined by three technical replicates. In the figure legends *n* is indicated and error bars indicate the standard error. Significant differences between samples were calculated using the student′s t-test and regarded as significant when P<0.05. Significant differences are indicated in the figures by asterisks (*P*<0.05 (*) or *P*<0.01 (**)).

